# The *hiuABC* Operon Mediates Xenosiderophore Utilization in *Caulobacter crescentus*

**DOI:** 10.1101/2025.09.04.674318

**Authors:** Sergio Hernandez-Ortiz, Aretha Fiebig, Sean Crosson

## Abstract

*Caulobacter* species are common residents of soil and aquatic ecosystems, where bioavailable iron is often extremely limited. Like other diderm bacteria, *C. crescentus* can acquire Fe(III) via outer-membrane TonB-dependent transporters (TBDTs) that recognize and import ferric siderophore complexes. Although *C. crescentus* is not known to synthesize siderophores, it encodes multiple TBDTs that are transcriptionally regulated by the ferric uptake repressor (Fur), suggesting it acquires iron by scavenging xenosiderophores produced by neighboring microbes. To identify *C. crescentus* genes required for xenosiderophore utilization, we developed a barcoded transposon screen using ferrioxamine B (FXB), a hydroxamate-family siderophore produced by soil actinomycetes, as a model substrate. This screen identified *hiuABC*, a conserved, Fur-regulated operon that supports FXB-dependent iron acquisition. We provide evidence that *hiuA* encodes the primary TBDT responsible for uptake of ferrioxamines and ferrichrome (FC), structurally distinct members of the hydroxamate siderophore family. *hiuB* encodes a PepSY-domain protein with structural similarity to *Pseudomonas aeruginosa* FoxB, a known periplasmic ferri-siderophore reductase. *hiuC* encodes a small, hypothetical membrane protein predicted to form a functional complex with HiuB in the inner membrane. Both *hiuB* and *hiuC* are required for utilization of FXB and ferrioxamine E (FXE), indicating a shared role in iron acquisition from ferrioxamines. Surprisingly, utilization of FC as an iron source required *hiuB* but not *hiuC*, suggesting a substrate-specific role for HiuC in ferri-siderophore processing. We conclude that the conserved *hiuABC* operon encodes a set of proteins that enable bacteria to acquire iron from structurally diverse hydroxamate-family siderophores.

**Importance:** Iron is often a limiting nutrient due to its poor solubility in the presence of oxygen. To overcome this, some microbes produce specialized molecules known as siderophores, which tightly bind and solubilize iron, facilitating its uptake into the cell. *Caulobacter* species are common in freshwater, marine, and soil environments, and there is emerging evidence they play important roles in plant-associated microbial communities. Here we report the discovery of a three-gene system that allows *C. crescentus* to acquire iron from a set of siderophores produced by select soil bacteria and fungi. We define functional roles for each protein component of this system, which informs a mechanism by which *Caulobacter* can pirate iron-scavenging molecules produced by its neighbors.

## Introduction

Iron is an essential nutrient required by nearly all organisms, serving as a cofactor in key biological processes including cellular respiration, DNA repair, and gas sensing (1). Though it is among the most abundant elements in the Earth’s crust (2), iron is often a limiting nutrient in oxygenated environments because ferric iron (Fe (III)) forms insoluble hydroxides and oxides at neutral pH (3). The concentration of bioavailable Fe (III) in aerobic soils and waters can be as low as 10^−18^ M at pH 7, far below the concentration needed to support microbial growth (4). In these conditions, microbes often obtain iron from their surroundings through the secretion of siderophores, which are high-affinity Fe (III) chelating molecules that solubilize iron from minerals or organic complexes (5). More than 500 siderophores have been identified across diverse ecological niches (6, 7).

Diderm bacteria face an additional challenge in iron acquisition due to their double-membrane cell envelope. Siderophores and other organic iron complexes are often too large or too dilute to rely on passive diffusion across the envelope. To actively acquire iron, diderms often employ TonB-dependent transporters (TBDTs), which are specialized β-barrel outer membrane receptors that bind and transport many substrates, including ferric-siderophore complexes (8). TBDTs exhibit specificity for particular structural classes of siderophores (e.g., catecholates, hydroxamates, or hydroxy-carboxylates) (9). Upon siderophore binding, the TBDT undergoes structural changes that expose its TonB box, enabling engagement with the TonB protein. TonB, which is energized via proton motive force from the ExbB–ExbD complex, drives structural rearrangements that actively import the ferri-siderophore into the periplasm (8). Once in the periplasm, ferri-siderophores may be routed to an ABC transporter for translocation into the cytoplasm or reduced by a periplasmic enzyme to liberate ferrous iron for import (10).

*Caulobacter* species are metabolically versatile (11–16) aerobic diderms that are common in terrestrial and aquatic ecosystems (17), and have been identified as hub species in plant microbial communities (18, 19). The well-studied model system, *Caulobacter crescentus*, lacks known siderophore biosynthesis genes but it encodes 62 predicted TBDTs. It is therefore likely that this bacterium has the capacity to recognize and import a diverse array of siderophores produced by other microorganisms (i.e., xenosiderophores) (20–22). Indeed, iron limitation is known to de-repress transcription of four Fur-regulated TBDT genes in *C. crescentus*: *cciT*, CCNA_00138, *hutA*, and CCNA_03023 (23). While *hutA* encodes a characterized heme/hemin transporter (20), the substrates for *cciT*, CCNA_00138, and CCNA_03023 (hereafter referred to as *hiuA*) remain undefined.

Hydroxamate siderophores including the linear trihydroxamate ferrioxamine B (FXB) and the cyclic hexapeptide ferrichrome (FC) are common in terrestrial ecosystems (7) and promote *C. crescentus* growth under iron-limiting conditions (20) Click or tap here to enter text.but the specific TBDTs mediating their uptake remain unidentified. Furthermore, the fate of ferric–siderophore complexes in the periplasm is poorly characterized in this important model bacterium. To address these gaps, we developed a broth-based genetic screening platform for analyzing siderophore acquisition mechanisms in *C. crescentus*. Here we report a genome-wide approach to identify siderophore acquisition genes in *Caulobacter crescentus* using a barcoded transposon mutant library. Building on prior work showing that EDTA treatment of complex medium induces iron starvation and that ferri-siderophore supplementation restores growth (24), we used FXB as a model exogenous iron source in our screen. Our approach uncovered a Fur-regulated three-gene ***<u>h</u>***ydroxamate ***i***ron ***<u>u</u>***ptake operon (*hiuA, hiuB, hiuC*) that supports utilization of structurally diverse hydroxamate-type ferri-siderophores that are common in soil and aquatic environments. This manuscript describes functional and structural analyses of the products of the *hiu* operon.

## Results

### Desferrioxamine limits intracellular iron and restricts *Caulobacter* growth, while FXB enhances growth under iron-limiting conditions

We began by testing whether 1 μM desferrioxamine B, the iron-free form of FXB, inhibits the growth of wild-type (WT) *C. crescentus* in peptone yeast extract (PYE) complex broth. Desferrioxamine B binds ferric iron with exceptionally high affinity (25) and thus will chelate and potentially sequester iron present in the growth medium. To assess its effect, we compared growth in the presence of 1 μM desferrioxamine B to that in 300 μM EDTA, a well-characterized cation chelator known to limit *C. crescentus* growth by reducing intracellular iron availability (24). The PYE broth produced in our laboratory contains approximately 700 nM iron (24), so the concentration of desferrioxamine used in this experiment exceeded available iron. The growth (OD_660_) of *C. crescentus* cultures was lower in the presence of desferrioxamine B than in untreated PYE or PYE treated with 300 μM EDTA (Figure 1A) indicating that *1)* desferrioxamine B is a more potent inhibitor of *C. crescentus* growth than EDTA, and *2)* the iron-complexed form of this siderophore (ferrioxamine B, or FXB) is not efficiently imported at the tested concentration.

**Figure 1.**
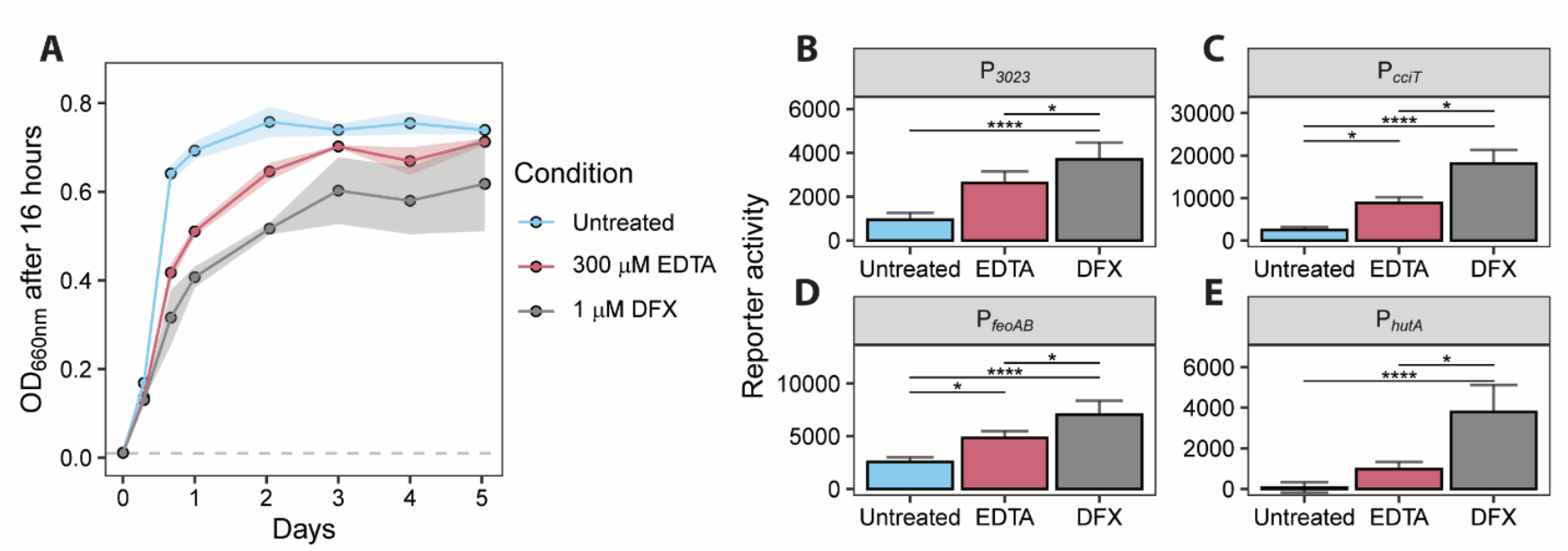
Growth inhibition by desferrioxamine is associated with reduced intracellular iron. **(A)** Optical density (OD_660_) of *C. crescentus* CB15 cultures grown in PYE broth for five days with or without 300 μM EDTA or 1 μM desferrioxamine B (DFX). Cultures were inoculated at an initial OD_660_ of 0.01 (dashed gray line). Values represent the mean ± SD (shaded areas) of six cultures grown in two independent experiments. **(B–E)** Fluorescence reporter assays measuring transcription from Fur-regulated promoters: **(B)** *feoAB* (PfeoAB), **(C)** *cciT* (PcciT), **(D)** *hutA* (PhutA), and **(E)** *hiuA* (PhiuA). Strains carrying promoter fusions to the mNeonGreen fluorescent protein were grown for 2 hours in PYE broth with or without 300 μM EDTA or 1 μM desferrioxamine. Bar plots show average fluorescence normalized to OD_660_, with error bars representing 1 standard deviation (n = 9). Statistical comparisons were performed using a Kruskal–Wallis test followed by Dunn’s post test. Significance: * *P* < 0.05; ****P < 0.0001.

We predicted that growth inhibition by desferrioxamine B was associated with intracellular iron limitation. To test this we measured transcription from four Fur-regulated promoters (P_*feoAB*_, P_*cciT*_, P_*hutA*_, P_*CCNA_03023*_) fused to a fluorescent reporter gene, mNeonGreen. Transcription from these Fur-regulated promoters was significantly higher in media containing 1 μM desferrioxamine B compared to untreated medium (Figure 1B) providing evidence that this concentration of desferrioxamine B reduces available iron in the cytoplasm, and thus attenuates *C. crescentus* growth via iron limitation. Moreover, the activity of each of these promoters was significantly higher in the presence of 1 μM desferrioxamine B than 300 μM EDTA providing evidence that 1 μM desferrioxamine B has a more potent iron limitation effect than 300 μM EDTA.

Finally, we tested whether addition of 1 μM FXB to EDTA-treated growth medium mitigated the growth-limiting effect of 300 μM EDTA. FXB binds ferric iron with an affinity 5–6 orders of magnitude greater than that of EDTA and is therefore expected to retain Fe(III) even in the presence of a 300-fold molar excess of EDTA (25). Growth of the WT strain was significantly improved with FXB supplementation relative to broth containing EDTA only, though the effect was modest (Figure S1). Evidence that this growth enhancement effect is due to acquisition of iron from FXB is presented below.

### A role for the CCNA_03023-21 operon (*hiuABC*) in iron acquisition from Ferrioxamine B

Our observation of growth enhancement by the ferri-siderophore FXB during EDTA chelation challenge (Figure S1), revealed an opportunity to identify genes that support iron acquisition from this siderophore. To this end, we used a Randomly Barcoded Transposon Sequencing (RB-TnSeq) method (26) in which we cultivated a *C. crescentus* Tn mutant library (27, 28) in PYE broth containing 300 μM EDTA, with or without 1 μM of Ferrioxamine B (FXB). Because FXB conferred only modest growth rescue in EDTA-treated broth, we serially passaged cultures in this experiment to amplify fitness defects (and possibly, advantages) associated with specific transposon insertions. This approach yielded mutant strains that appeared to be defective in iron acquisition from FXB (Figure 2; Table S1).

**Figure 2.**
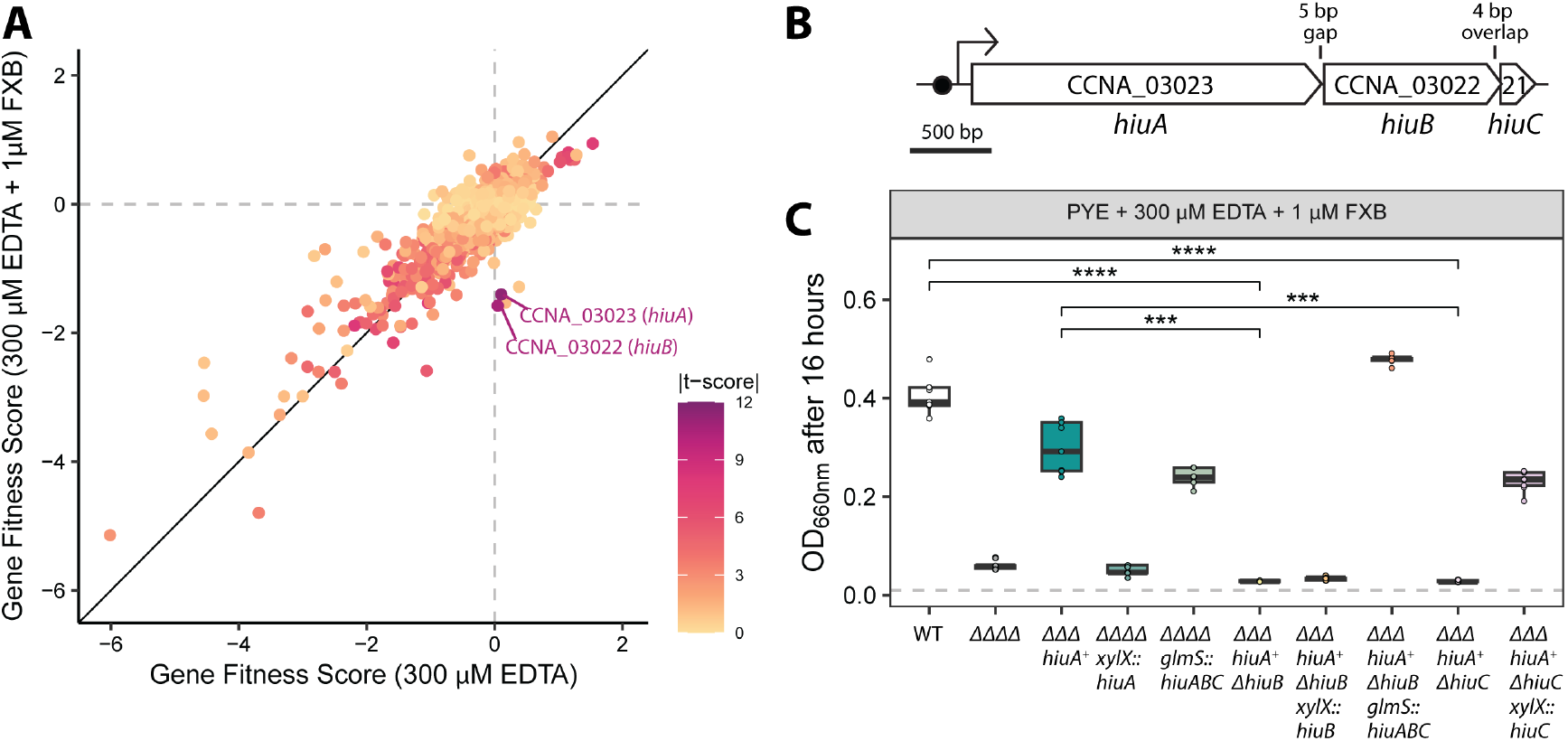
Genetic screen identifies genes in the *hiu* operon that support *C. crescentus* fitness during growth on ferrioxamine B (FXB) as the sole iron source. **(A)** Gene fitness scores derived from a genome-wide screen using a barcoded *C. crescentus* Tn-himar mutant library after growth in complex PYE medium supplemented with 300 µM EDTA (iron-depleted condition) or with 300 µM EDTA plus 1 μM FXB. Each point represents the mean fitness score for a single gene, averaged across four independent cultures per condition. Dot color reflects the *t*-score associated with each fitness value, providing a measure of statistical confidence in the fitness eeect. Genes required specifically for growth in the presence of FXB exhibit no fitness defects in the presence of only EDTA but negative fitness scores when grown with FXB. Complete fitness scores are provided in Table S1. **(B)** Diagram of the *hiuA–hiuB–hiuC* locus (*CCNA_03023–03021*). A predicted Fur-binding site (Fur box) upstream of the locus is indicated by a black circle. **(C)** Optical density (OD_660_) measurements of *C. crescentus* strains grown for 16 hours in PYE broth supplemented with 300 µM EDTA and 1 µM ferrioxamine B (FXB). Strains are wild-type (WT), lacking all four fur-regulated TonB-dependent transporters including hiuA (*ΔΔΔΔ*), lacking 3 fur-regulated TonB-dependent transporters and encoding only *hiuA* (*hiuA*^+^ΔΔΔ), and derived strains lacking either *hiuB* or *hiuC* and/or carrying ectopic expression constructs for *hiuA, hiuB, or hiuC*. Full strain genotypes can be found in Table S2. Cultures were inoculated at an initial OD_660_ of 0.01 (dashed gray line) and grown for 16 hours. Box plots show the median and interquartile range (25th–75th percentiles), overlaid with individual replicate values (n = 9). Comparisons of OD_660_ between strains were performed using a Kruskal-Wallis followed by Dunn’s post test. Statistical significance is indicated as follows: ***P < 0.001, ****P < 0.0001.

Inspection of mutant fitness scores identified three gene categories (Figure 2A): (a) mutants with no detectable fitness change in any condition; (b) mutants with similar fitness defects in both EDTA treated and EDTA plus FXB-supplemented media, and (c) mutants with fitness defects specifically in the FXB-supplemented condition but not in media containing only EDTA. This third group included strains harboring insertions genes including CCNA_00973 (encoding a protein tyrosine phosphatase), CCNA_03022 (a PepSY-family inner membrane protein), CCNA_03023 (a TonB-dependent outer membrane transporter, TBDT), and CCNA_03277 (a glycosyltransferase).

To prioritize candidates for further analysis, we inspected the t-scores associated with mutant fitness values under FXB-supplemented conditions. These scores reflect the statistical confidence of fitness defects (26). CCNA_03022 and CCNA_03023 mutants had the highest t-scores (t >10), indicating that strains harboring insertions in these genes had consistent and significant growth defects. CCNA_03023 encodes a TBDT related to *E. coli* FhuA (20), a transporter known to mediate the uptake of ferric hydroxamate siderophores (29). CCNA_03022 encodes a predicted inner membrane protein of the PepSY-PiuB superfamily (30). These two genes, along with CCNA_03021 (encoding a predicted 61-residue hypothetical inner membrane protein) comprise a Fur-regulated operon (23, 31) (Figure 2B). CCNA_03021 was not identified in our RB-TnSeq screen as it contains only a single TnHimar (i.e. TA dinucleotide) insertion site, though this gene is syntenic with CCNA_03022 and CCNA_03023 across Proteobacteria and in Bacteriodota (Figure S2) suggesting it shares a functional role in iron acquisition from FXB. We hereafter refer to these genes as hydroxamate iron uptake A through C: *hiuA* (gene locus CCNA_03023), *hiuB* (gene locus CCNA_03022) and *hiuC* (gene locus CCNA_03021).

### HiuA is the primary TonB-dependent transporter for ferrioxamine B uptake, and its function requires HiuB and HiuC

To test whether *hiuA* encodes the primary TonB-dependent transporter (TBDT) for ferrioxamine B (FXB) uptake, we used a set of strains in which three of the four Fur-regulated TBDTs have been deleted and thus this set of strains individually encode only one Fur-regulated TBDTs: HiuA (*ΔΔΔ hiuA+*), CciT (*ΔΔΔ cciT+*), CCNA_00138 (*ΔΔΔ 138+*), or HutA (*ΔΔΔ hutA+*). In PYE medium containing 300 μM EDTA, addition of 1 μM FXB had a marginal effect (∼10% increase) on the growth of WT and *ΔΔΔ cciT*^+^ strains (Figure S3 and S4). The *ΔΔΔ 138*^+^ and *ΔΔΔ hutA*^+^ strains failed to grow in the presence of EDTA and showed a modest FXB-dependent growth improvement (Figures S3 and S4). In contrast, the *hiuA*^+^ strain exhibited a ∼200% increase in growth in response to FXB addition. We conclude that HiuA is the primary outer membrane transporter for FXB in *C. crescentus*.

To dissect the functional contribution of other genes in the *hiuABC* operon to FXB-dependent growth enhancement, we generated in-frame deletions of *hiuB* and *hiuC* in the *ΔΔΔ hiuA*^+^ background, yielding *ΔΔΔ hiuA*^+^ *ΔhiuB* and *ΔΔΔ hiuA*^+^ *ΔhiuC* strains. A strain lacking all four Fur-regulated TBDTs (ΔΔΔΔ) served as a negative control. In the presence of EDTA and FXB, the *ΔΔΔ hiuA*^+^ strain exhibited enhanced growth, while the ΔΔΔΔ, *ΔΔΔ hiuA*^+^ *ΔhiuB*, and *ΔΔΔ hiuA*^+^ *ΔhiuC* strains failed to show FXB-dependent growth enhancement (Figure 2C). Ectopic expression of *hiuC* alone (driven by the native *hiuABC* promoter) restored FXB-dependent growth in the *ΔΔΔ hiuA*^+^ *ΔhiuC* strain, but ectopic expression of *hiuA* or *hiuB* individually from the native promoter did not restore growth in the corresponding deletion strains. However, expression of the full *hiuABC* operon successfully complemented both *ΔΔΔΔ* and *ΔΔΔ hiuA*^+^ *ΔhiuB* strains (Figure 2C). Notably, complementation of *ΔΔΔ hiuA*^+^ *ΔhiuB* with the full *hiuABC* operon restored growth to WT levels, which is higher than the parent *hiuA*^+^ background. This result suggests that coordinated expression of all three genes facilitates more efficient FXB uptake, perhaps by optimizing stoichiometry or enhancing protein levels. Altogether, these results provide evidence that HiuA serves as the primary FXB transporter in *C. crescentus*, and that acquisition of iron from FXB (transported via HiuA) requires the inner membrane proteins, HiuB and HiuC.

### The inner membrane proteins HiuB and HiuC support FXB utilization independently of the outer membrane import route

The ΔΔΔΔ strain lacks all four Fur-regulated TBDTs (including HiuA) but exhibited a slight growth enhancement when supplemented with 1 μM FXB (Figure 3), suggesting the existence of alternative, lower-affinity FXB acquisition mechanisms. To test this, we cultured WT, *ΔΔΔ hiuA*^*+*^, and ΔΔΔΔ strains in PYE + 300 μM EDTA that was supplemented with increasing concentrations of FXB (1– 10 μM). FXB enhanced growth in all strains in a concentration-dependent manner, with the *hiuA*^*+*^ strain reaching WT-like levels of growth at 3 μM FXB (Figure 3A). Growth of the ΔΔΔΔ strain was impaired relative to WT, as expected, but was still enhanced by increasing FXB. We conclude that ferric iron from FXB can be acquired via mechanisms that do not require any of the four known Fur-regulated TBDTs.

**Figure 3.**
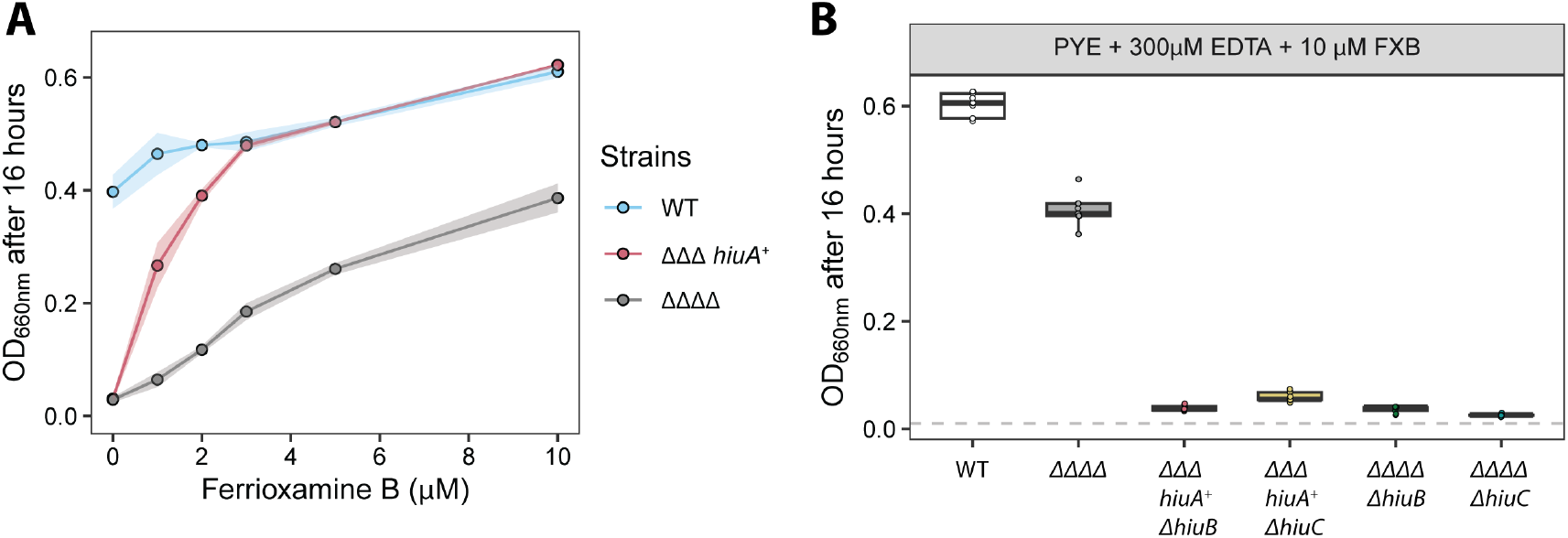
Utilization of ferrioxamine B (FXB) requires *hiuA, hiuB* and *hiuC*. **(A)** Growth (OD_660_) of wild-type (WT), ΔΔΔ *hiuA*^+^, and the ΔΔΔΔ strain (lacking all four fur-regulated TonB-dependent transporters, including *hiuA*), grown for 16 hours in PYE medium containing 300 µM EDTA and increasing concentrations of FXB. Each point represents the mean and shaded areas represent the standard deviation of nine biological replicates performed on separate days. **(B)** Endpoint OD_660_ values of WT, ΔΔΔΔ, and mutants lacking *hiuB* or *hiuC* in the ΔΔΔ *hiuA*^*+*^ and ΔΔΔΔ backgrounds cultures after 16 hours of growth in PYE with 300 µM EDTA and 10 µM FXB. Box plots show the median and interquartile range (25th–75th percentiles), overlaid with individual data points (n = 9). Cultures were inoculated at OD_660_ = 0.01 (dashed gray line).

To test whether the inner membrane proteins HiuB and HiuC function specifically with the outer membrane transporter HiuA or are more broadly required for FXB utilization, we measured the growth of *ΔΔΔ hiuA*^*+*^ *or ΔΔΔΔ* strains in which *ΔhiuB* or *ΔhiuC* were deleted in PYE + 300 μM EDTA supplemented with 10 μM FXB (i.e. the concentration that produced the greatest growth enhancement in the ΔΔΔΔ strain). Neither *ΔΔΔ hiuA+ ΔhiuB* nor *ΔΔΔ hiuA+ ΔhiuC* grew under these conditions (Figure 3B). Similarly, deleting either *hiuB* or *hiuC* in the ΔΔΔΔ background eliminated the FXB-dependent growth enhancement observed at high FXB concentrations. We conclude that *hiuA* provides the most efficient route of FXB import across the outer membrane, but that *hiuB* and *hiuC* are required to process the iron from FXB regardless of the outer membrane entry route.

### The *hiuABC* operon supports iron acquisition from ferrioxamine E

Since the *hiuABC* operon supports FXB-dependent growth enhancement during EDTA chelation challenge, we tested whether it also supports iron acquisition from the related (but structurally distinct) cyclic trihydroxamate siderophore, ferrioxamine E (FXE). Supplementation of PYE broth + 300 μM EDTA with 1 μM FXE did not increase culture density after 16 hours (Figure S5), which indicated that *C. crescentus* either lacks an efficient uptake mechanism for FXE or simply cannot acquire iron from FXE.

To distinguish between these possibilities, we increased the FXE concentration to 10 μM. The ΔΔΔΔ strain lacking all four Fur-regulated TBDTs showed a modest growth improvement at 10 μM FXE, while the *ΔΔΔ hiuA+* strain exhibited a more pronounced growth rescue (Figure S5). As with FXB, FXE-dependent growth enhancement required both *hiuB* and *hiuC*. These results indicate that iron acquisition from FXE, although less efficient than FXB, also proceeds through mechanism that requires the HiuB and HiuC inner membrane proteins.

### HiuA and HiuB, but not HiuC, are required for ferrichrome utilization

Our results provided evidence that the *hiuABC* operon facilitates iron acquisition from the structurally related trihydroxamate siderophores FXB and FXE. To test whether *hiuABC* also supports utilization of hydroxamate siderophores belonging to other structural classes, we measured *C. crescentus* growth in PYE medium containing 300 μM EDTA supplemented with 1 μM ferrichrome (FC), a cyclic hexapeptide hydroxamate.

Growth of the ΔΔΔΔ strain, which lacks all four Fur-regulated TBDTs, was only weakly enhanced by ferrichrome (FC), suggesting minimal acquisition of iron from FC in this background (Figures 4 and 5). In contrast, the strain encoding only *hiuA* (*ΔΔΔ hiuA*^+^) exhibited wild-type levels of growth in the presence of 1 μM FC, indicating that HiuA enables efficient import of FC. Deletion of *hiuB* in this background (*ΔΔΔ hiuA*^+^ *ΔhiuB*) abolished the growth enhancement conferred by FC, while deletion of *hiuC* (*ΔΔΔ hiuA*^+^ *ΔhiuC*) had no effect (Figure 4). These results support a model in which HiuB is necessary for iron acquisition from FC transported through the HiuA outer membrane receptor, while HiuC is not. Genetic complementation of *hiuA*^+^ *ΔhiuB* with ectopically-expressed *hiuB* restored FC-dependent growth, as did expression of the full *hiuABC* operon. Similarly, ectopic expression of *hiuA* in the ΔΔΔΔ background was sufficient to restore FC-dependent growth confirming a role for *hiuA* and *hiuB* in ferrichrome utilization.

**Figure 4.**
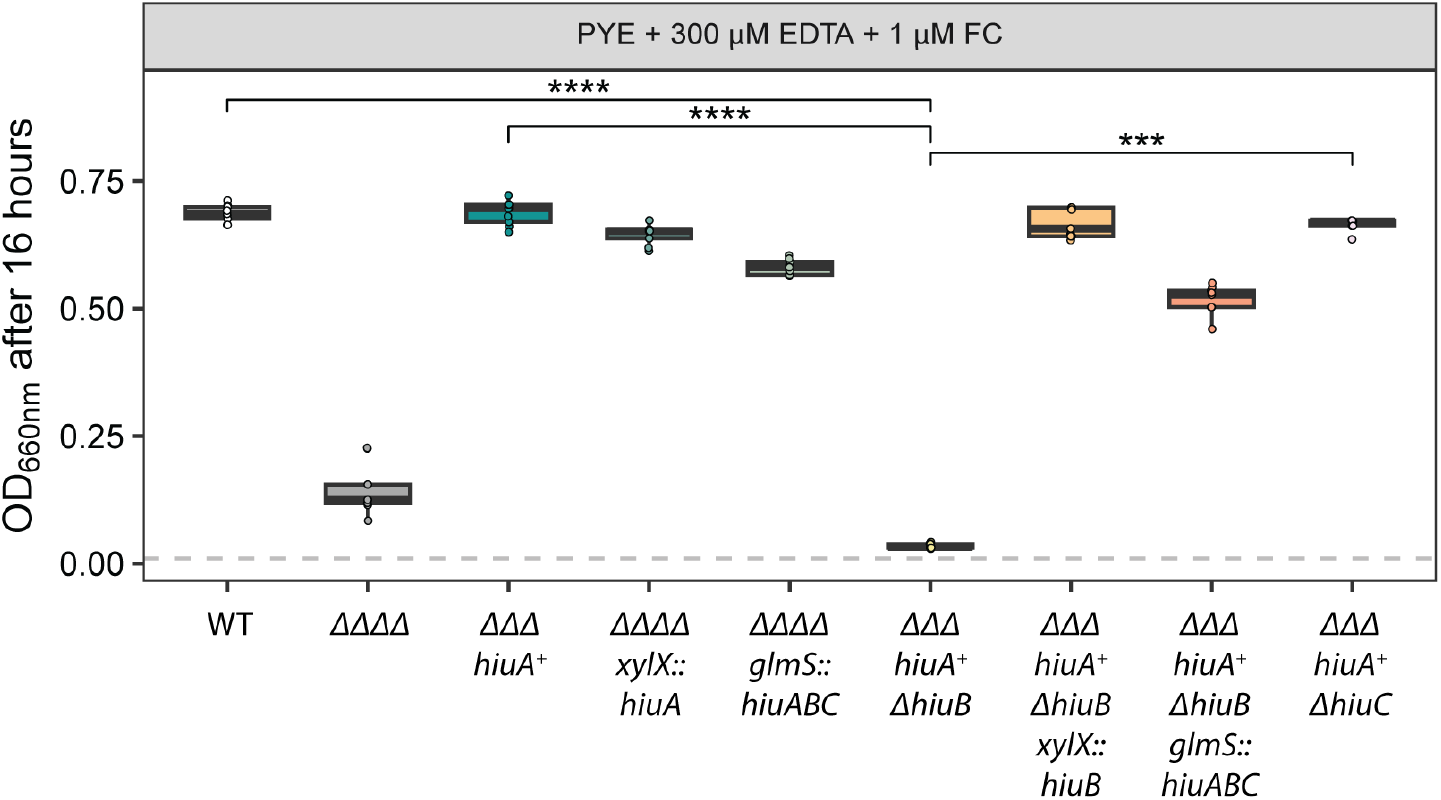
HiuA and HiuB support *C. crescentus* growth on ferrichrome. Optical density (OD_660_) measurements of *C. crescentus* strains grown for 16 hours in PYE broth supplemented with 300 µM EDTA and 1 µM ferrichrome (FC). Strains include wild-type (WT), ΔΔΔΔ (lacking all four Fur-regulated TBDTs), *hiuA*^+^ΔΔΔ (encoding only *hiuA*), strains derived from *hiuA*^+^ΔΔΔ with additional deletions in either *hiuB* or *hiuC*, and strains complemented with the respective deleted genes. Box plots show the median, 25th, and 75th percentiles, overlaid with individual data points from each independent culture (n = 9). Statistical comparisons of OD_660_ between strains were performed using a Kruskal–Wallis test followed by Dunn’s post test. Significance is indicated as: ***P < 0.001, ****P < 0.0001.

**Figure 5.**
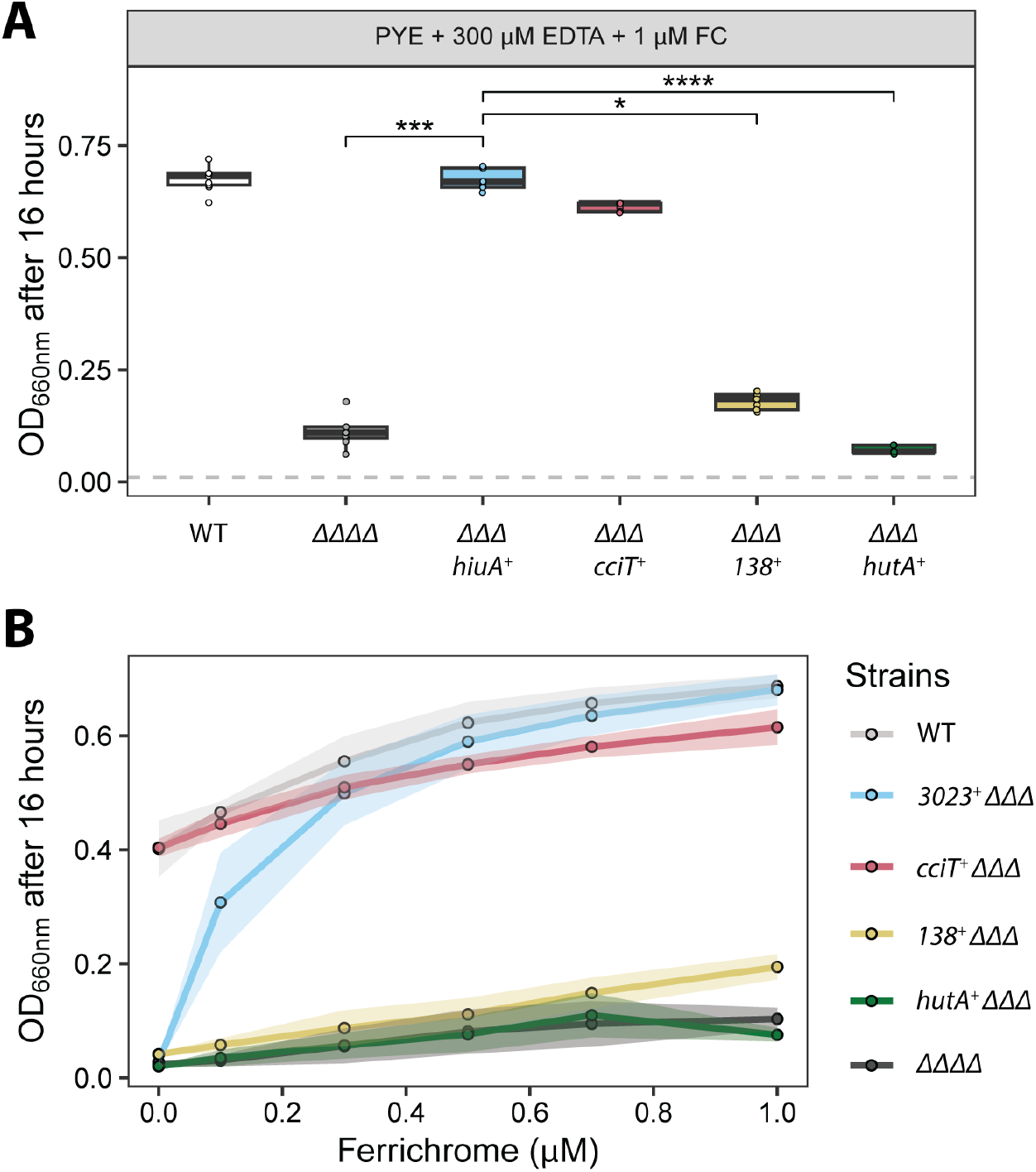
HiuA is the primary outer membrane importer of ferrichrome in *C. crescentus*. **(A)** Optical densities (OD_660_) of strains after 16 hours of growth in PYE broth treated with 300 μM EDTA and supplemented with 1 μM Ferrichrome. Strains include wild-type (WT), a strain lacking all four Fur-regulated TBDTs *(ΔΔΔΔ*), and strains encoding only one of each of the four Fur-regulated TonB-dependent transporters (*cciT, CCNA_00138, hutA, or hiuA*) while lacking the other three (*ΔΔΔ*). Cultures were inoculated at OD_660_ = 0.01 (dashed gray line). Box plots display the median and interquartile range (25th–75th percentiles), overlaid with individual data points from independent cultures (n = 9). Statistical comparisons of OD_660_ values were performed using a Kruskal-Wallis test followed by Dunn’s multiple comparisons test. Significance levels are denoted as follows: P < 0.05 (*) and P < 0.0001 (****). **(B)** Optical densities (OD_660_) of strains after 16 hours of growth in PYE broth treated with 300 μM EDTA and supplemented with an increasing concentration of Ferrichrome. Each point represents the mean of nine independent replicates; shading indicates 1 standard deviation.

We next compared FC-dependent growth enhancement across strains encoding single Fur-regulated TBDTs. As previously shown, *cciT* confers wild-type-like growth in the presence of EDTA (24), while *hiuA* alone enabled wild-type growth in the presence of 1 μM FC and uniquely supported growth at FC concentrations as low as 100 nM (Figure 5). In contrast, strains encoding only *CCNA_00138* or *hutA* exhibited only modest growth at 1 μM FC and failed to grow appreciably at lower FC levels (Figure 5B). These results indicate that HiuA functions as a high-affinity FC transporter and suggest it is the only Fur-regulated TBDT capable of supporting efficient iron acquisition from this cyclic peptide hydroxamate. We conclude that acquisition of iron from low-concentration FC requires both HiuA and HiuB, but not HiuC.

## Discussion

Siderophore-mediated iron acquisition is a widespread bacterial strategy for overcoming iron limitation in aerobic environments (32). In diderms, this process requires active transport across the outer membrane via TonB-dependent transporters (TBDTs), which specifically bind siderophores within defined structural classes (8). Once in the periplasm, ferri-siderophores can be transported to the cytoplasm by inner membrane transporters or reduced to liberate ferrous iron for subsequent import across the inner membrane. *C. crescentus* does not encode known siderophore biosynthesis genes, but possesses 62 predicted TonB-dependent transporters (TBDTs), several of which are transcriptionally regulated by the ferric uptake repressor (Fur) (20, 22, 33). This suggests a growth/survival strategy centered on scavenging xenosiderophores produced by neighboring organisms in the environment.

We report that the Fur-regulated (34) *hiuABC* operon, composed of three broadly co-conserved genes (Fig. S2), encodes proteins that collectively support the utilization of multiple hydroxamate siderophores. HiuA is a TBDT that is related (20) FhuA, the FC receptor of *E. coli*. We have provided evidence that HiuA is the primary outer membrane receptor for FXB, FXE, and FC. The inner membrane proteins HiuB (a PepSY–PiuB superfamily protein) and HiuC (a small uncharacterized hypothetical protein) are both required for FXB and FXE utilization, regardless of the outer membrane import route. Surprisingly, these proteins differed in their requirement for ferrichrome utilization: *hiuB* was necessary for FC utilization, while *hiuC* was not. Thus it appears that HiuC has a substrate-specific role in ferri-siderophore processing.

Structural modeling with AlphaFold3 (35) predicts a high-confidence interaction (ipTM = 0.90) between HiuB and HiuC in the inner membrane (Figure 6A), supporting the hypothesis that these proteins function together as a complex. HiuB contains PepSY domains, which have been associated with periplasmic reductase activity in other systems (36). If the HiuB-HiuC complex catalyzes siderophore iron reduction in the periplasm, this would explain the requirement of both genes for utilization of ferrioxamine siderophores. Supporting this hypothesis, AlphaFold3 modeling of HiuB reveals two putative histidine-ligated heme-binding sites within its transmembrane domain (Figures 6A and 6B). These correspond to experimentally resolved heme ligation sites in the structure of the *P. aeruginosa* ferrioxamine reductase FoxB, which are proposed to mediate electron transfer to ferrioxamine via a through-space tunneling mechanism (30). We propose that HiuC modulates the reductase activity of HiuB, potentially by influencing substrate specificity. This model aligns with our observation that *hiuC* is required for utilization of ferrioxamines (FXB and FXE) but dispensable for ferrichrome (FC), suggesting substrate-dependent regulation of HiuB activity by HiuC.

**Figure 1.**
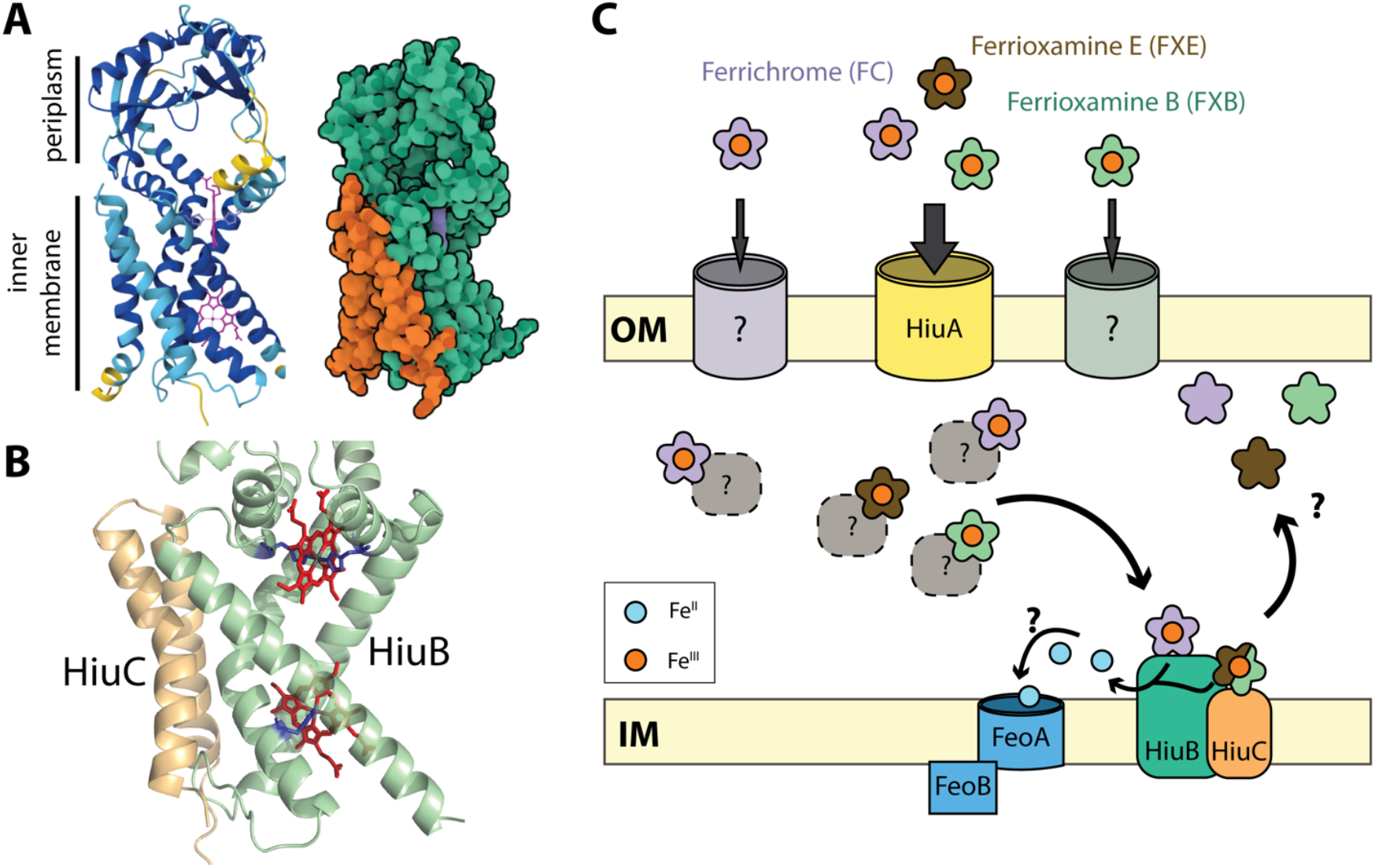
HiuABC-dependent utilization of hydroxamate siderophores in *C. crescentus*. **(A)** *left-*Ribbon diagram structural model of the predicted HiuB:HiuC complex with inner membrane and periplasmic domains labeled. Ribbon cartoon is colored by Alphafold3 pLDDT prediction confidence score from yellow (low) to dark blue (high). Two heme cofactors are shown in purple (ipTM=0.90; pTM=0.88). *right*-Space-filling model of the predicted HiuB:HiuC complex in the same orientation as left. HiuB is shown in green; HiuC in orange. **(B)** Zoomed in view of the transmembrane region of proteins HiuB (green) and HiuC (yellow) showing the predicted heme (red) binding sites. The conserved histidine residues that ligate the heme iron center as in PDB:7ABW are shown in dark blue. The lower heme cofactor has only one predicted histidine ligation site, while the upper heme cofactor has two. The bound heme cofactors in HiuB are shown in red. **(C)** Schematic model of ferri-siderophore uptake are periplasmic Fe(III) reduction mediated by the HiuABC system in *C. crescentus*. At the outer membrane (OM), HiuA mediates high-affinity import of FXB, FXE and FC. There is evidence that FC also enters through a route independent of the Fur-regulated TBDTs (*huiA, cciT, hutA* and *CCNA_00138*). In the periplasm, imported siderophores may be bound by yet to be identified periplasmic binding proteins (dotted line-gray circles). At the inner membrane (IM), HiuB (a PepSY-family protein) and HiuC (a small inner membrane protein) are required for FXB and FXE utilization, possibly by facilitating reductive iron release from the siderophore. In contrast, ferrichrome utilization requires HiuB, but not HiuC. The FeoAB system is proposed to mediate import of ferrous iron (Fe(II)) into the cytoplasm following reduction.

The *hiuABC* system appears to be mechanistically distinct from the well-studied *fhu* ferrichrome uptake system in *E. coli* (29, 37). In the *fhu* system, the outer membrane TBDT (FhuA) is functionally linked to a periplasmic binding protein (FhuD) and an inner membrane ABC transporter (FhuB-FhuC). In contrast, *hiuABC* operon lacks ABC transporter components; the route of iron or ferri-siderophore transport across the inner membrane remains unknown. It seems likely that HiuB (and in some cases an HiuB-HiuC complex) promotes reductive release of hydroxamate-bound iron in the periplasm, thereby facilitating transfer of ferrous iron to an as-yet-unidentified transporter. The FeoAB ABC transport system (38) is a likely candidate for this ferrous iron transport role (Figure 6C).

Our data also support the existence of low-affinity, HiuA-independent uptake routes for FXB and FC. This functional redundancy aligns with the expansive repertoire of TBDTs in *C. crescentus* and is consistent with result from other bacteria where select substrates can be transported by multiple TBDTs (8, 39). Notably, however, HiuA was the only Fur-regulated TBDT tested that enabled high-affinity uptake of FC at environmentally relevant concentrations (100 nM), consistent with its role as the primary (and likely physiological) transporter of these hydroxamate siderophores under iron-limiting conditions typical of soil and aquatic environments. Ferrioxamines and ferrichrome are produced by diverse microbes and are commonly found in terrestrial and aquatic ecosystems (6, 7, 40). The ability of *C. crescentus* to utilize these hydroxamates via the *hiuABC* system likely enhances its fitness in such iron-scarce habitats. The conserved gene synteny of *hiuABC* across diverse bacterial phyla, including Proteobacteria and Bacteroidota (Fig. S2), suggests that this operon functions broadly as a ferri-hydroxamate utilization module in many genera.

We conclude by noting that the broth-based, genome-wide screening approach reported here enables direct linkage of specific TBDTs to their ferri-siderophore substrates. In the future, this scalable, quantitative platform can facilitate identification of other iron acquisition systems. Creation of a barcoded *C. crescentus* library in a background lacking the major (and somewhat promiscuous) *cciO–cciT* iron import system (24) could improve dynamic range of this screening approach and reveal TBDT-ferri-siderophore pairs that provide only a weak growth advantage in the presence of a competing iron acquisition system. Beyond siderophores, and as shown previously (41), this approach can be adapted to investigate uptake of other metals or organic nutrients in defined or natural media (27) providing new insights into bacterial strategies for nutrient acquisition across ecosystems.

## Materials and Methods

### Growth conditions and strain construction

*C. crescentus* CB15 and derived strains were grown at 30°C in peptone-yeast extract (PYE) medium (0.2% [w/v] peptone (Fisher Bioreagents, Lot No. 225155), 0.1% [w/v] yeast extract (Fisher Bioreagents, Lot No. 220635), 1 mM MgSO_4_ (Fisher Chemical, Lot No. 183674), 0.5 mM CaCl_2_ (Fisher Chemical, Lot No. 117031)) and is best described as a complex medium. The Fe•EDTA chelate (Sigma-Aldrich, F0518, Lot No. RNBD1641) used to supplement PYE, is a 1:1 molar mix of FeSO_4_ and EDTA (ferrous sulfate chelate solution kept at 4°C, Sigma-Aldrich, F0518). When ferric iron was used, a 100 mM FeCl_3_ aqueous stock (pH ∼3.0) kept at −20°C was thawed and used to make a working 10 mM stock that was used to generate the ferri-siderophores by mixing 1:1 molar ratio to reach the final concentrations indicated. When an apo- or iron-bound siderophore (desferrichrome/Ferrichrome (*Ustilago sphaerogena*; Sigma-Aldrich), desferrioxamine B/Ferrioxamine B, Ferrioxamine E (*Streptomyces antibioticus*, Millipore) was added to liquid media, it was added to sterile media just before bacterial inoculation. The water used for the preparation of all media was ultra-purified using a Barnsted GenPure water purification system (ThermoFisher). *E. coli* strains were grown at 37°C in LB medium (1% [w/v] tryptone, 0.5% [w/v] yeast extract, 1% [w/v] NaCl). Growth media were solidified by the addition of 1.5% (w/v) agar when necessary. To grow any strain lacking *cciT* on a PYE solid medium, the medium was supplemented with 10 μM Fe•EDTA (Sigma-Aldrich, F0518). Antibiotics were used at the following concentrations in liquid and solid media, respectively, as appropriate: *C. crescentus*, 5 µg/mL or 25 µg/mL kanamycin, 1 µg/mL or 2 µg/mL chloramphenicol, 1 µg/mL or 2 µg/ml tetracycline, 20 µg/mL nalidixic acid; *E. coli*, 50 µg/mL kanamycin, 20 µg/mL chloramphenicol, 10 µg/mL tetracycline.

Standard molecular biology techniques were used to construct all plasmids. Detailed information on strains, plasmids, and primers can be found in Table S2. To create plasmids for in-frame deletion allele replacements, the regions flanking the target gene were cloned into pNPTS138 (42). For the generation of fluorescent transcriptional reporter plasmids, promoter regions (200–500 bp upstream of the open reading frame) were cloned into the vector pPTM056 (43). For extopic expression of individual genes, the entire coding sequences plus stop codon (with their respective promoter regions) were cloned into pMT585 (pXGFPC2) (13), which integrates at the *xylX* locus. In the case of *hiuB* and *hiuC*, the genes were amplified and stitched together with the *hiuABC* operonic promoter before being cloned into pMT585. Ectopic expression of the operon was achieved by amplifying and cloning the entire coding sequence and promoter into a mini-TN7 cassette in pUC18-mTn7K. The cassette was then integrated at the glmS locus using a helper plasmid, pTNS3, encoding the TN7 transposase. All plasmids were introduced into *C. crescentus* by conjugation. For allele replacements, a double recombination strategy was used to select merodiploid strains harboring each pNPTS-based plasmid with kanamycin resistance, followed by *sacB* counter selection on PYE plates containing 3% (w/v) sucrose. PCR was used to evaluate colonies that were sucrose-resistant and kanamycin-sensitive to identify those colonies harboring the null allele. For the construction of strains with multiple TonB-dependent transporter gene deletions, counterselection was performed on PYE sucrose plates supplemented with 10 µM Fe•EDTA (Sigma-Aldrich, F0518).

### Tn-himar-seq to assess genes that support growth enhancement by Ferrioxamine B (FXB) in the presence of EDTA

An aliquot of a *C. crescentus* CB15 barcoded Tn-himar mutant library (27, 28, 44) was grown in 5 mL of liquid PYE medium containing 5 μg/mL kanamycin for 8 hours. One milliliter of this starter culture was set aside as a reference, and for each condition (PYE, PYE with 300 μM EDTA, and PYE with 300 μM EDTA and 1 μM Ferrioxamine B) four replicate 2 mL cultures were inoculated to a starting optical density at 660 nm (OD_660_) of 0.01. Cultures were grown in 100 × 14 mm glass tubes, shaking at 200 RPM at 30°C. After 16 hours, cultures were back diluted to an OD_660_ of 0.01 and incubated in the same conditions for an addition 16 hours for a total of 32 hours of growth. After incubation, 1 mL of each culture was collected by centrifugation at 16,000 × g for 3 minutes. The supernatants were discarded, and the resulting cell pellets were resuspended in 10–20 μL of water and stored at –20°C.

Barcode abundances were determined following the workflow developed by Wetmore et al (26). Briefly, barcodes were amplified using Q5 polymerase (New England Biolabs) in a 20-μL reaction containing Q5 reaction buffer, GC enhancer, 0.8 units Q5 polymerase, 0.2 mM dNTPs, 0.5 μM of each primer, and 1 μL of the cell suspension, using the following amplification conditions: 98°C for 4 min, 25 cycles of (98°C for 30 sec, 55°C for 30 sec, and 72°C for 30 sec), 72°C for 5 min, followed by a hold at 4°C. The primer set consisted of a universal forward primer, Barseq_P1, and a uniquely indexed reverse primer, Barseq_P2_ITxxx, with the index number denoted by xxx (see ref (26)). The barcode amplification products were pooled and sequenced as 50-bp single-end reads on an Illumina NovaSeq instrument, using Illumina TruSeq primers. Barcode sequence analysis was carried out using the fitness calculation protocol of Wetmore et al (26). Barcodes from each sample were counted and compiled using MultiCodes.pl and combineBarSeq.pl scripts. A barcode table was generated, from which the FEBA.R script was used to calculate fitness score relative to the reference starter culture. Scripts were provided by Morgan Price and are available at https://bitbucket.org/berkeleylab/feba/src/master/.

### Growth measurements

A single colony from a strain grown on a PYE plate was inoculated into 2 mL of liquid PYE medium and grown overnight. The next day, the cultures were diluted to an OD_660_ of 0.15 and outgrown for 2 h before being used to inoculate fresh cultures in the desired experimental condition (medium) at an OD_660_ of 0.01. Cultures were then incubated at 30°C while shaking at 200 rpm. After 16 h of growth, the optical density of the cultures was measured. For extended growth experiments (Figure 1), optical density was measured at 7, 16, 24, 49, 72, 96, & 121 hours.

### Analysis of transcription using fluorescent reporters

Strains harboring fluorescent transcriptional reporter plasmids to monitor the activity of the *feoB cciT, hutA or hiuABC* promoters (fused to mNeonGreen) were cultured overnight in triplicate at 30°C in liquid PYE containing 1 µg/mL chloramphenicol. These starter cultures were diluted to an OD_660_ of 0.001 and incubated overnight at 30°C with shaking at 200 rpm. The next morning, these cultures were used to inoculate 500 µL of liquid medium in the wells of a clear 48-well plate. The plate lids were sealed onto the plate body with AeraSeal sealing film (RPI Research Products International) and incubated at 30°C while shaking at 155 RPM. After 2 hours, OD_660_ and fluorescence (excitation at 497 ± 10 nm; emission at 523 ± 10 nm) were measured on a Tecan Spark 20M plate reader, and fluorescence values were normalized to the corresponding optical density. Statistical analysis was conducted using R.

## Acknowledgements

Research reported in this publication was supported in part by the National Institute of General Medical Science award R35GM131762 and by Army Research Office contract W911NF2210105 to S.C.

---

**Figure S1.**
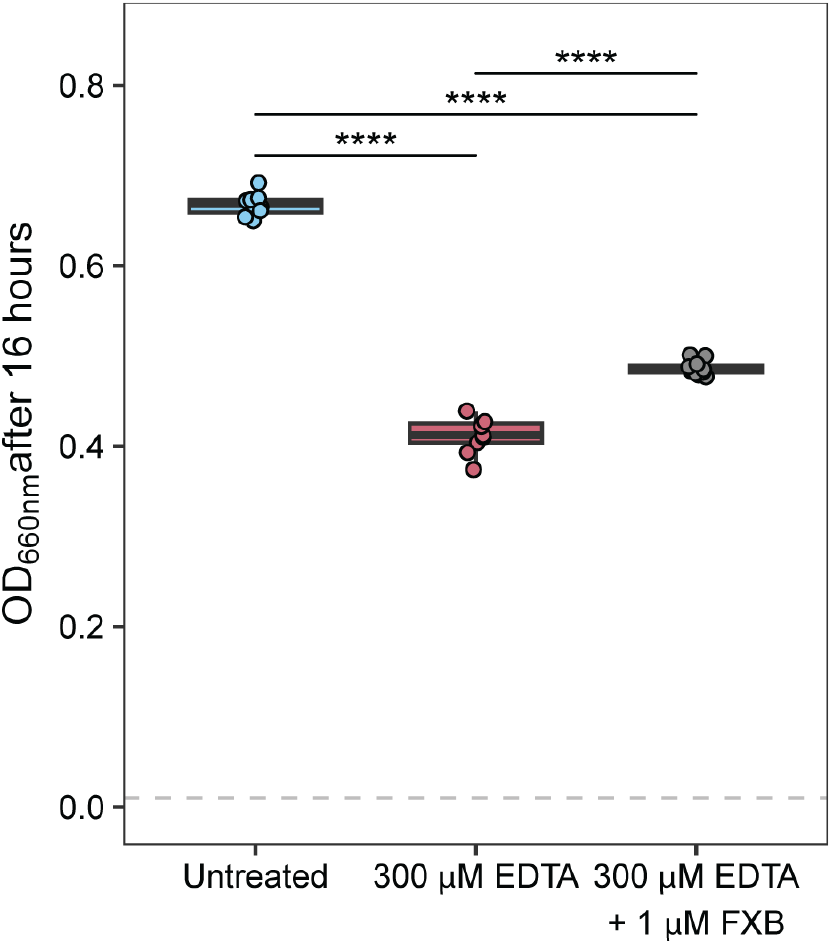
Ferrioxamine B enhances the growth yield of *C. crescentus* in iron-limited PYE broth. Optical density (OD_660_) of strains after 16 hours of growth in PYE broth under the following conditions: untreated (blue), supplemented with 300 μM EDTA (pink), or with 300 μM EDTA and 1 μM ferrioxamine B (FXB). Box plots show the median and interquartile range (25th and 75th percentiles), overlaid with individual data points for each independent culture (n = 9). Statistical comparisons of OD_660_ values were performed using a Kruskal–Wallis test followed by Dunn’s post hoc test. Significance: ****P < 0.0001.

**Figure S2.**
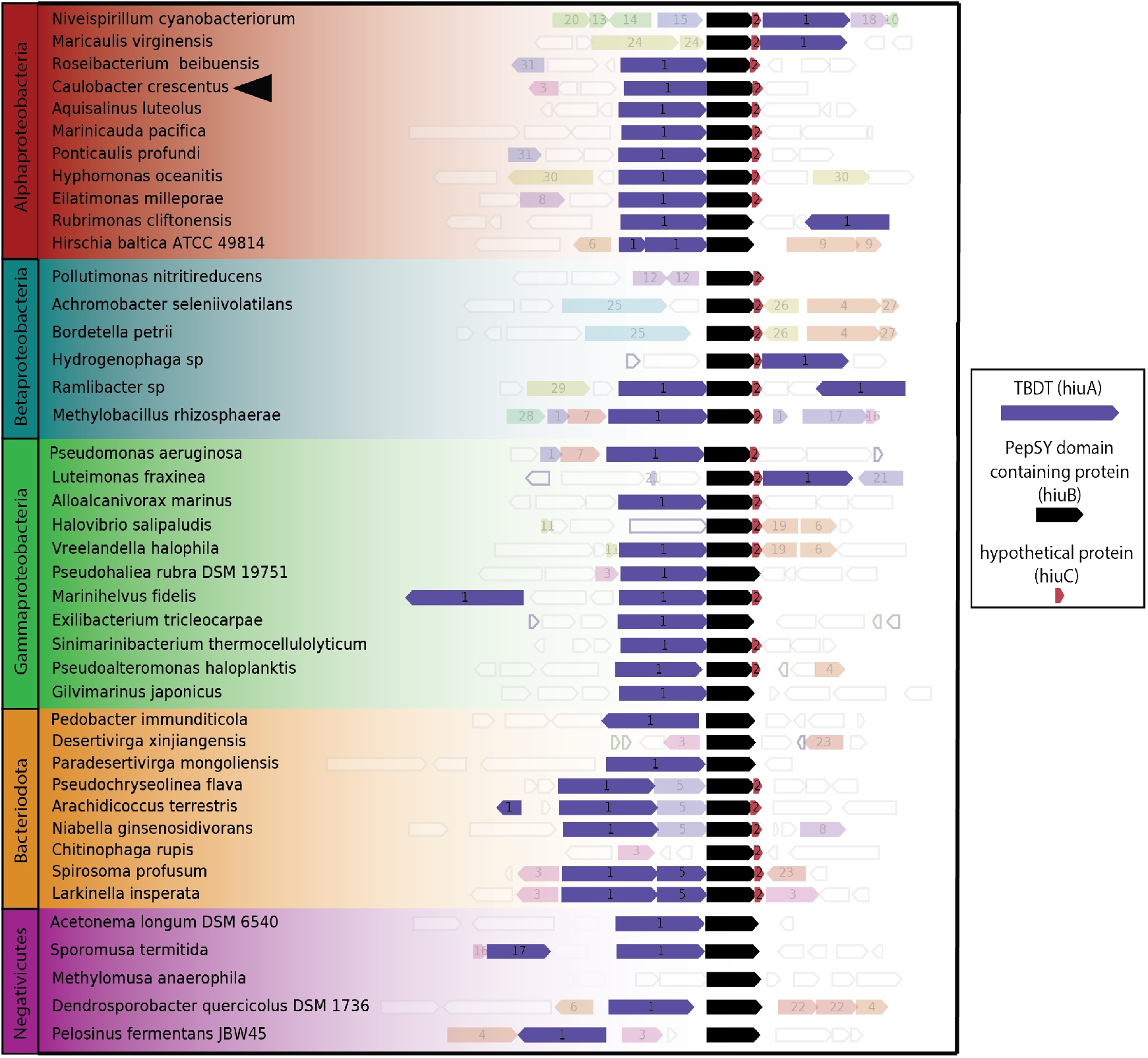
Conserved hiuABC-like operons are found across diverse bacterial lineages. Gene neighborhood analysis centered on *hiuB* reveals conserved synteny of *hiuA, hiuB*, and *hiuC* orthologs across multiple bacterial classes. Representative neighborhoods were selected based on the highest-scoring BLAST hits from members of the Proteobacteria, Bacteroidota, and Bacillota. Gene neighborhoods were visualized using webFLAGs (45). The *Caulobacter crescentus* locus is indicated by a black arrow. Orthologs of *hiuA, hiuB*, and *hiuC* are colored in purple, black, and red, respectively.

**Figure S3.**
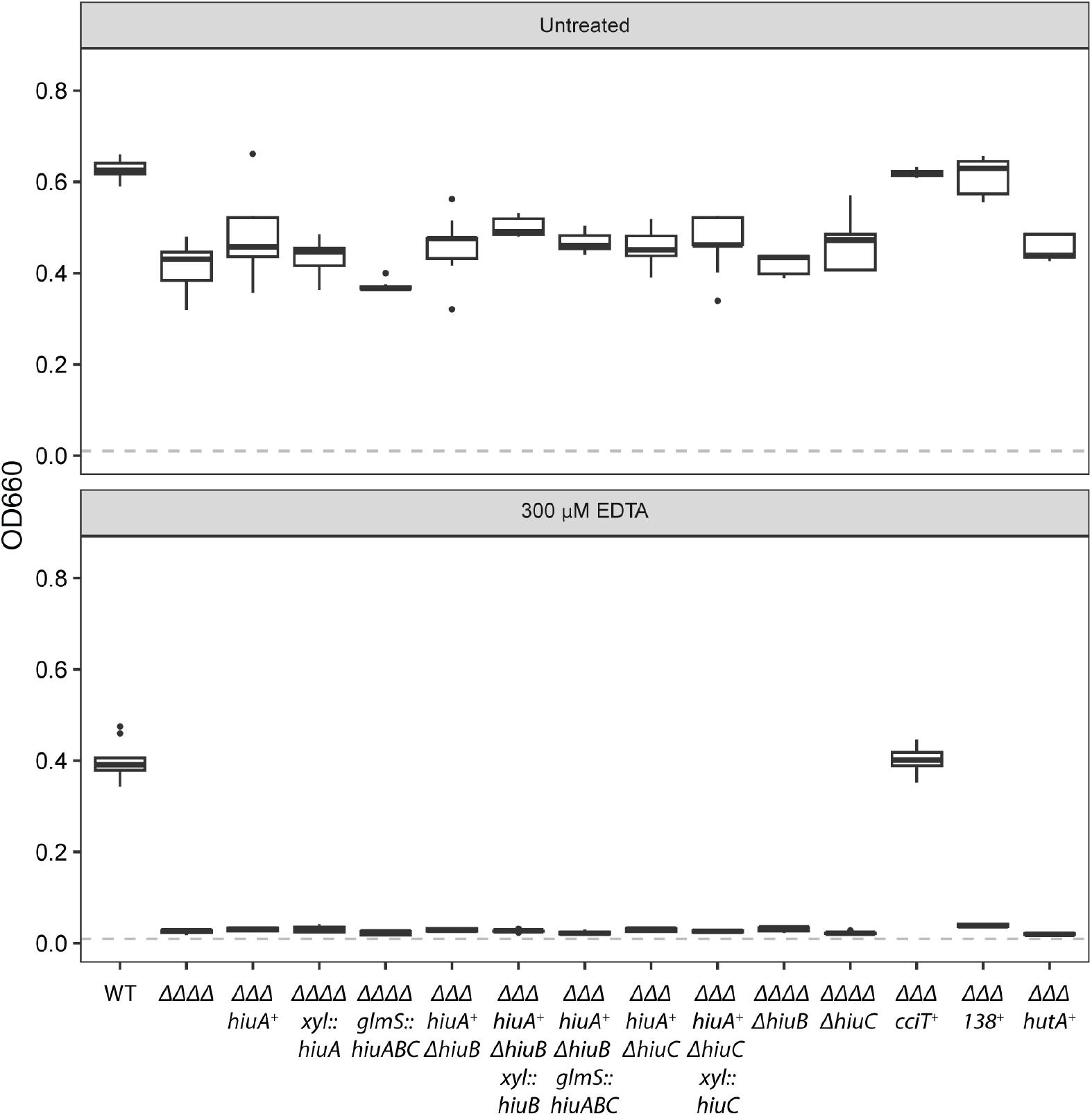
Growth of all *C. crescentus* strains in untreated and EDTA-treated PYE broth. Optical density at 660 nm (OD_660_) was measured after 16 hours of growth for all *C. crescentus* strains used in this study. Full genotypes and corresponding abbreviated genotypes for all strains are included in Table S2. Cultures were inoculated at an initial OD_660_ of 0.01 (indicated by the dashed gray line) and grown in either standard PYE broth or PYE broth treated with 300 µM EDTA (B). Box plots display the median and interquartile range (25th to 75th percentiles) and are overlaid with individual data points from nine independent cultures per strain (n = 9).

**Figure S4.**
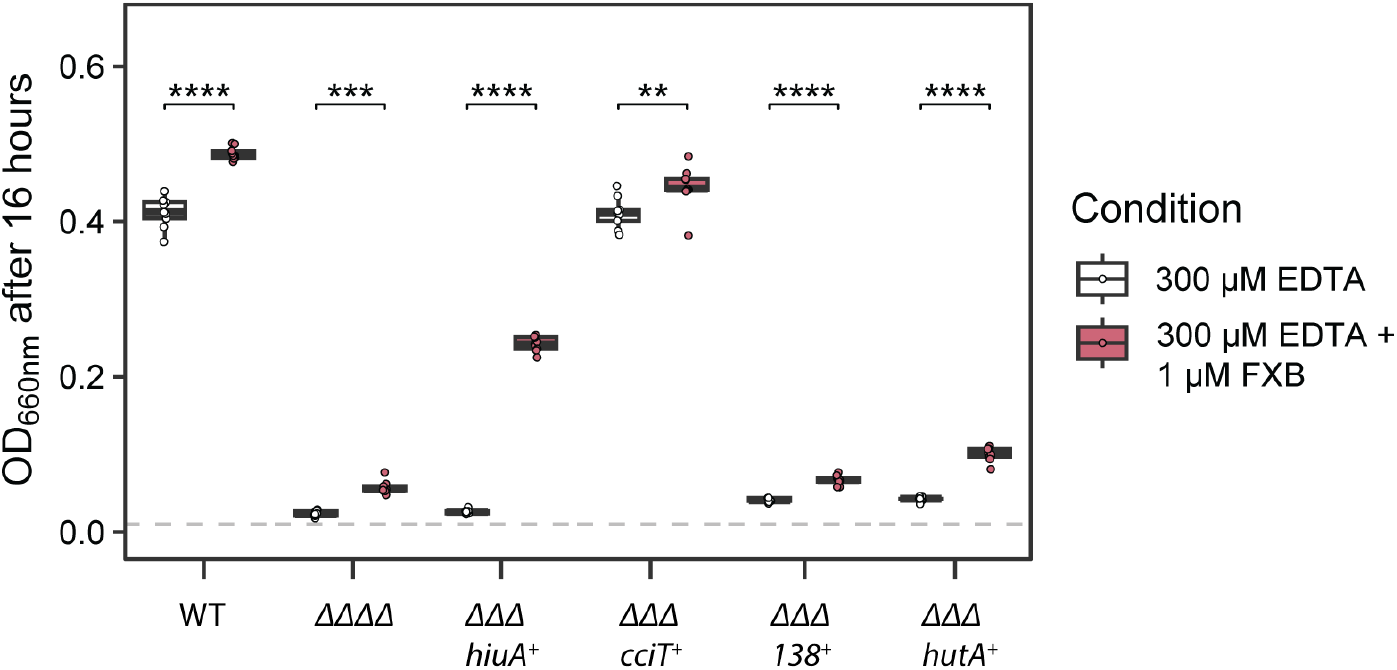
*hiuA* confers optimal growth on Ferrioxamine B. Optical density (OD) at 660 nm of cultures growth 16 hours in PYE broth treated with 300 μM EDTA and supplemented with 1 μM Ferrioxamine B (FXB). Strains include wild-type (WT), a strain lacking all four Fur-regulated TBDTs *(ΔΔΔΔ*), and strains encoding only one of each of the four Fur-regulated TBDTs (*cciT, CCNA_00138, hutA, or hiuA*) and lacking the other three (*ΔΔΔ*) Fur-regulated TBDTs. Strains were inoculated at 0.01 OD represented by the dashed gray line. Box plots reflecting the median and the 25th and 75th percentiles are overlaid with the individual data points for each independent culture (n = 9). Comparison of the OD of a strain between conditions was done through an unpaired t-test (Wilcoxon rank-sum test for *cciT*^*+*^). Statistical significance is indicated as follows: **P < 0.01, ***P < 0.001, ****P < 0.0001.

**Figure S5.**
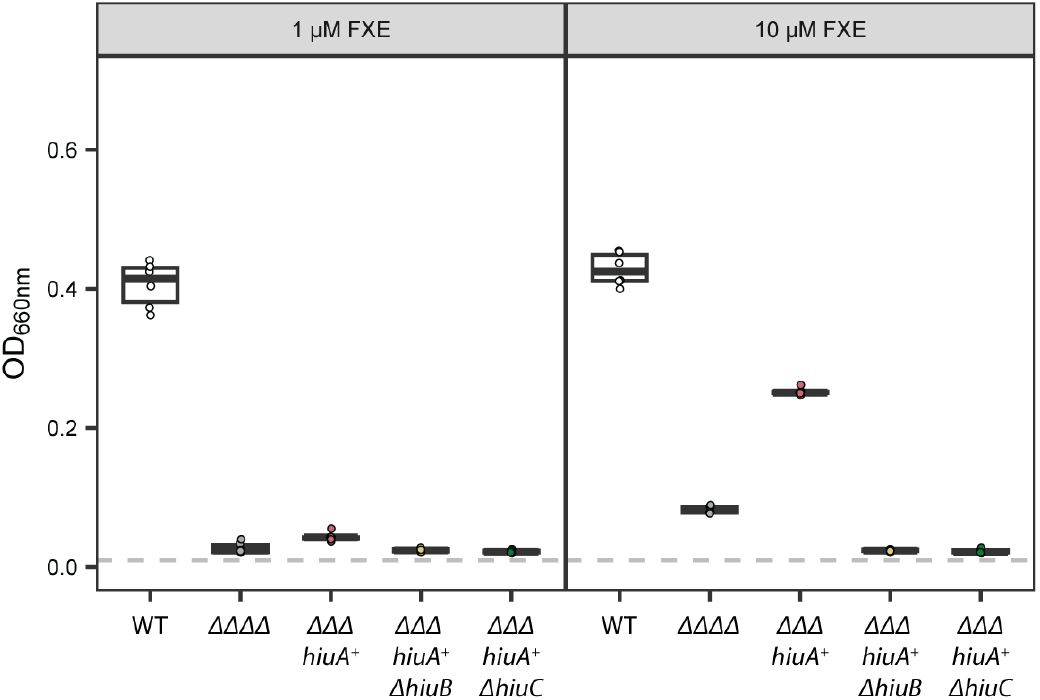
The *hiuABC* operon supports growth on Ferrioxamine E. Optical density (OD_660_) of cultures grown in PYE with 300 µM EDTA and 1 µM Ferrioxamine E (FXE) or 10 µM FXE broth for 16 hours. Strains are wild-type (WT), the ΔΔΔΔ strain (lacking all four fur-regulated TonB-dependent transporters, including *hiuA*), *hiuA*^+^*ΔΔΔ* (encoding only *hiuA*), strains lacking either *hiuB*, or *hiuC* in the *hiuA*^+^ *ΔΔΔ* background. Strains were inoculated at 0.01 OD660 represented by the dashed gray line. Box plots reflecting the median and the 25th and 75th percentiles are overlaid with the individual data points for each independent culture (n =9).

